# Genomic selection for lentil breeding: empirical evidence

**DOI:** 10.1101/608406

**Authors:** Teketel A. Haile, Taryn Heidecker, Derek Wright, Sandesh Neupane, Larissa Ramsay, Albert Vandenberg, Kirstin E. Bett

## Abstract

Genomic selection (GS) is a type of marker-based selection which was initially suggested for livestock breeding and is being encouraged for crop breeding. Several statistical models and approaches have been developed to implement GS; however, none of these methods have been tested for use in lentil breeding. This study was conducted to evaluate different GS models and prediction scenarios based on empirical data and to make recommendations for designing genomic selection strategies for lentil breeding. We evaluated nine single-trait models, two multiple-trait models, and models that account for population structure and genotype-by-environment interaction (GEI) using a lentil diversity panel and two recombinant inbred lines (RIL) populations that were genotyped using a custom exome capture assay. Within-population, across-population and across-environment predictions were made for five phenology traits. Prediction accuracy varied among the evaluated models, populations, prediction scenarios, traits, and statistical models. Single-trait models showed similar accuracy for each trait in the absence of large effect QTL but BayesB outperformed all models when there were QTL with relatively large effects. Models that accounted for GEI and multiple-trait (MT) models increased prediction accuracy for a low heritability trait by up to 66% and 14% but accuracy did not improve for traits of high heritability. Moderate to high accuracies were obtained for within-population and across-environment predictions but across-population prediction accuracy was very low. This suggests that GS can be implemented in lentil to make predictions within populations and across environments, but across-population prediction should not be considered when the population size is small.

## Introduction

Lentil (*Lens culinaris* Medik.) is an important pulse crop and a dietary source of protein, carbohydrate, micronutrients, vitamins, and fiber. Lentil, along with other pulses, is used increasingly as an important source of protein for people whose diet is solely plant-based. Moreover, it has ecological advantages as a rotational crop in cereal-based cropping systems by enabling better management of pests, herbicide residue, and soil nitrogen. In 2016, lentil was grown on over 5.5 Mha across 52 countries, accounting for an annual production of 6.3 million tonnes (FAOSTAT 2018). Until recently, however, lentil has received little attention in terms of genetic research. Advances in high-throughput genotyping technologies and a relative decline in sequencing costs have resulted in the availability of abundant single nucleotide polymorphisms (SNPs) covering the gene space of lentil through in-depth sequencing of the entire set of exons in the genome (Ogutcen et al. 2018). This new development has led to the availability of dense genetic maps and opened up opportunities to evaluate genomic tools such as genomic selection (GS) for lentil improvement.

Genomic selection is a type of marker-based selection that uses a training population (TP) that is phenotyped and genotyped to train a statistical model which is then used to predict genomic estimated breeding values (GEBVs) of individuals in a breeding population that have been genotyped but not phenotyped (Meuwissen et al. 2001; Meuwissen 2009). The GEBVs are then used for the selection of superior lines rather than the actual phenotypes. Model prediction accuracy is commonly tested using independent lines in a validation population that also has phenotypic and genotypic data. This is commonly achieved by partitioning the same population into training and validation sets. Genomic estimated breeding values are predicted for individuals in the validation set and the correlation of predicted values with the actual phenotypes is considered as the prediction accuracy.

Unlike the standard marker assisted selection (MAS) which uses a small number of markers associated with major QTL, GS uses a large sets of genome-spanning markers to potentially capture all the QTL underlying a trait (Meuwissen et al. 2001). Fitting all markers simultaneously avoids multiple testing and the need to identify markers-trait associations based on an arbitrarily chosen significance threshold. Genomic selection was initially suggested for livestock breeding (Meuwissen et al. 2001), and later evaluated and implemented in crop breeding, especially for wheat and maize (Bernardo 2009; Bernardo and Yu 2007; Beyene et al. 2015; Crossa et al. 2016; Crossa et al. 2014). The goal of genomic selection is to enhance the genetic gain of quantitative traits by accelerating the breeding cycle and increasing selection intensity (Battenfield et al. 2016; Heffner et al. 2010). Thus far, there is no information on the utility of genomic selection for lentil breeding.

The accuracy of GS is affected by several factors, including TP size, marker density, trait heritability, genetic relationship between the training and breeding populations, linkage disequilibrium (LD), population structure, GEI, and the statistical model used (Burgueño et al. 2012; Calus 2010; Combs and Bernardo 2013; de los Campos et al. 2015; Heffner et al. 2011; Jarquín et al. 2014; VanRaden et al. 2009; Wientjes et al. 2013). A large TP size, higher heritability, and higher marker density usually improve genomic prediction accuracy (Asoro et al. 2011; Heffner et al. 2011; Liu et al. 2018; Meuwissen et al. 2001). However, even marker distribution across the genome is essential to tag important QTL and maintain GS accuracy (Bassi et al. 2016). The main assumption of GS is that each QTL is in LD with at least one nearby marker and potentially all the genetic variance can be explained by the markers (Calus 2010; Goddard and Hayes 2007). It has also been shown that incorporating GEI into GS models can improve the prediction accuracy (Crossa et al. 2015; de los Campos et al. 2015; Jarquín et al. 2017; Lopez-Cruz et al. 2015). The effect of population structure on prediction accuracy varies based on prediction strategies, genetic architectures of traits, and the type of population (Guo et al. 2014). Therefore, it is important to evaluate models that account for GEI and population structure in each population and breeding environment.

Genomic selection models fit a large number of markers on a small number of phenotypic observations which may lead to overfitting when markers are treated as fixed effects (de los Campos et al. 2013). Standard GS models treat markers as a random effect and reduce the dimension of the marker data either by selecting variables, shrinking marker effect estimates or a combination of both (de los Campos et al. 2013). Several methods, including ridge regression, Bayesian regression, kernel-based approaches, and machine learning algorithms have been developed that differ mainly in their assumptions about the variances of marker effects (Breiman 2001; Gianola et al. 2006; Gianola and van Kaam 2008; Habier et al. 2011; Meuwissen et al. 2001; Park and Casella 2008; Pérez et al. 2010). Ridge regression assumes that all loci explain equal amount of the genetic variance, while Bayesian models allow the variance to vary across loci (Meuwissen et al. 2001). The assumption of common variance for all loci in a ridge regression is unrealistic and may underestimate the effects of known major QTL. Alternatively, models that treat markers associated with traits, markers tagging candidate genes, and previously discovered QTL, as fixed effects have been developed (Spindel et al. 2016). Initially, GS models were developed for predicting a single trait from a single environment. In recent years, multivariate models and models that account for GEI have been developed (Burgueño et al. 2012; Jarquín et al. 2014; Jia and Jannink 2012; Jiang et al. 2015; Lopez-Cruz et al. 2015). Multivariate and multi-environment models utilize the correlation of traits and environments to enhance GS accuracy (Jia and Jannink 2012; Lopez-Cruz et al. 2015). To date, none of these models have been tested in lentil; thus, the objectives of this study were to evaluate different GS models and prediction scenarios based on empirical data in lentil and to make recommendations for designing genomic selection strategies for lentil breeding.

## Materials and Methods

### Plant material and phenotyping

We used three populations to evaluate genomic prediction accuracy in lentil. The first population was a lentil diversity panel (LDP) composed of 324 accessions obtained from the gene banks of the International Center for Agricultural Research in the Dry Areas (ICARDA), United States Department of Agriculture (USDA), Plant Gene Resources of Canada (PGRC), as well as cultivars developed at the Crop Development Centre (CDC), University of Saskatchewan (http://knowpulse.usask.ca/portal/project/AGILE%3A-Application-of-Genomic-Innovation-in-the-Lentil-Economy; click on germplasm to see the full list). These lines were evaluated at Sutherland (lat 52°09’, long 106°30’) from 2016 to 2018 and Rosthern (lat 52°41’, long 106°17’), Saskatchewan, in 2016 and 2017. The second population, LR-01, was composed of 110 recombinant inbred lines (RILs) developed from a cross between ‘CDC Robin’ and ‘ILL 1704’. ‘CDC Robin’ is a high yielding, red cotyledon lentil cultivar developed by the CDC, University of Saskatchewan (Vandenberg et al. 2002). ‘ILL 1704’ is a landrace from Ethiopia. The third population, LR-11, included 120 RILs developed from a cross between ‘CDC Milestone’ and ‘ILL 8006-BM4’. ‘CDC Milestone’ is a high yielding, yellow cotyledon lentil cultivar developed at the CDC (Vandenberg et al. 2001). ‘ILL 8006-BM4’ was derived from ‘Barimasur-4’, a high-yielding, disease-resistant, red cotyledon cultivar from Bangladesh (Sarker et al. 1999). The LR-11 population was evaluated at Sutherland and Rosthern, SK, in 2017 and 2018. LR-01 was grown at the same two locations in 2018. For all populations, field trials were established in 1-m^2^ micro-plots with three seeded rows. A seeding rate of 60, 100, and 120 seeds per m^2^ were used for the LDP, LR-01, and LR-11, respectively. The field experiments were arranged in a randomized complete block design with three replications in each site-year. Plots were seeded from late April to mid-May and harvested from mid-August to early September in each year.

Phenotypic traits, including days to flowering (DTF), vegetative period (VEG), days to swollen pods (DTS), days to maturity (DTM), and reproductive period (REP), were measured in all populations. Days to flowering and DTS were recorded when 10% of plants in a plot had at least one open flower and pods with fully swollen seeds, respectively. Days to maturity was taken when 10% of plants in a plot had half of their pods mature. Vegetative and reproductive periods were recorded as the number of days from emergence to flowering and from flowering to maturity, respectively. The phenotypic data were analyzed using analysis of variance (ANOVA) with SAS Mixed models, v9.4 (SAS Institute Inc. 2015). For each population, the phenotypic data were analyzed separately in each environment (site-year) and combined across environments. Genotype (i.e. lines) was considered as a fixed effect and replication was considered random for the analysis of data in each environment. For combined analysis of data, replication nested in environment, environment (site-years), and GEI were considered random effects. The Kenward-Roger degrees of freedom approximation method was used to compute the degrees of freedom for means (Kenward and Roger 1997). Broad-sense heritability (H^2^) was estimated in each population using the equation 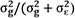, where 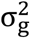 and 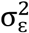 are the genetic and residual variances calculated using the ‘lmer’ function in the R package lme4, v1.1-7 (Bates et al. 2016).

### Genotypic data

Genomic DNA was extracted from fresh leaves of two to three-week-old seedlings for all three populations. For each line, DNA was extracted from leaves of one to two plants using a Qiagen MiniPrep Kit (Qiagen). Genotyping of all lines was performed using a custom lentil exome capture assay as described in Ogutcen et al. (2018). Markers with more than 5% missing data and a minor allele frequency of less than 5% were removed prior to analysis. A total of 9394, 24,395, and 39,297 markers that were common between the LDP and LR-11, LDP and LR-01, and LR-01 and LR-11, respectively, were used for across-population genomic prediction. Within-population genomic predictions were made using the 24,395 markers in the LDP and 39,297 markers in LR-01 and LR-11. Missing marker genotypes were imputed using the function ‘A.mat’ in R package rrBLUP, v4.4 (Endelman 2011).

### Statistical models and prediction scenarios

#### A model that accounts for population structure

The effect of population structure on model prediction accuracy was assessed in the LDP using an interaction model that accommodates genetic heterogeneity by partitioning marker effects into components that are common across groups and random deviations that are group specific as described in de los Campos et al. (2015). Marker-based K-means clustering was applied to partition the 324 lines into three groups. There were 77, 106, and 141 lines in groups one, two, and three, respectively. The interaction model assumes that marker effects have two components, one that is common across groups (*b*_0*k*_ where *k* = 1, 2, … *p* refers to markers) and another that is group specific (*b*_1*k*_, *b*_2*k*_ and *b*_3*k*_ for groups 1, 2 and 3, respectively). The marker effects are *β*_1*k*_ = *b*_0*k*_ + *b*_1*k*_, *β*_2*k*_ = *b*_0*k*_ + *b*_2*k*_, and *β*_3*k*_ = *b*_0*k*_ + *b*_3*k*_ for groups one, two, and three, respectively. The equation of the interaction model for the three groups becomes

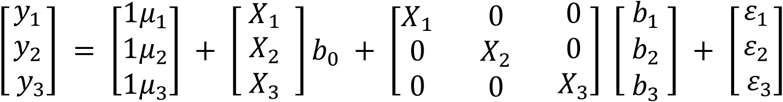

where *y*_1_, *y*_2_, *y*_3_, and *X*_1_, *X*_2_, *X*_3_ represent the phenotypes and genotypes of the individuals in groups one, two, and three, respectively, *μ*_1_, *μ*_2_, and *μ*_3_ are group specific intercepts, *b*_0_, *b*_1_, *b*_2_, and *b*_3_ are vectors of marker effects and *ε*_1_, *ε*_2_, and *ε*_3_ represent model residuals. Prediction accuracy of the interaction model was compared with an across-group model that assumes constant marker effects across groups, thereby ignoring population structure. The across-group model can be obtained by setting *b*_1_ = *b*_2_ = *b*_3_ = 0; therefore, the regression equation for the three groups becomes

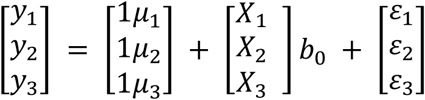

The interaction and across-group approaches were fitted using BayesB model in R (R Development Core Team 2016), using the Bayesian generalized linear regression (BGLR) package, v1.0.8, (Pérez and de los Campos 2014). Prediction was made using 10 random training-validation partitions. In each partition, 260 lines (62 in group one, 85 in group two and 113 in group three) were used as TP and 64 lines (15 in group one, 21 in group two and 28 in group three) were used for cross-validation. Prediction accuracy was obtained by correlating within-group predicted values with phenotypes of lines in the validation set.

#### Standard single-trait prediction models

Single-trait predictions were made using ridge regression best linear unbiased prediction (RR-BLUP), genomic best linear unbiased prediction (G-BLUP), BayesA, BayesB, BayesCπ, Bayesian Lasso (BL), Bayesian ridge regression (BRR), and Bayesian reproducing kernel Hilbert spaces regression (RKHS). The RKHS regression is a semi-parametric approach which accounts for both additive and non-additive genetic effects (de los Campos et al. 2009), while all the other models are based only on the additive effect. These models were fitted in R (R Development Core Team 2016), using the BGLR package, v1.0.8, (Pérez and de los Campos 2014), and the ridge regression and other kernels for genomic selection (rrBLUP) package, v4.4 (Endelman 2011). The default settings of BGLR (five degrees of freedom and the scale parameter based on sample variance of the phenotypes) were used (Pérez and de los Campos 2014). For G-BLUP, the genomic relationship matrix (GRM) was computed according to VanRaden (2008). For RKHS, we used a Gaussian kernel with a bandwidth parameter of *h* = 1/M × {1/5, 1, 5} as described in Pérez and de los Campos (2014), where M is the median squared Euclidean distance between all lines calculated using off-diagonals.

#### A model that incorporates the results of genomewide association studies

We used a GS+GWAS model to fit significant markers identified from fold specific GWAS as fixed effects in GS (Spindel et al. 2016). GS+GWAS is equivalent to RR-BLUP when no marker is fitted as a fixed effect (Spindel et al. 2016). The accuracy of GS+GWAS was tested using a five-fold cross-validation design. In the LDP, GWAS was conducted in each fold based on the phenotypic and genotypic data of the TP using compressed mixed linear model (CMLM) in GAPIT, v3.0 (Lipka et al. 2012). Population structure and cryptic relatedness were accounted for using five marker-derived principal components and kinship as covariates. In LR-01 and LR-11, single marker regression was performed in each fold based on the phenotypic and genotypic data of the TP using the ‘lm’ function in R (R Development Core Team 2016). For all three populations, the *P*-values were sorted from low to high and multiple testing correction was performed based on a False Discovery Rate (FDR) using the function “p.adjust” and “BH” method in R (R Development Core Team 2016). Then the SNPs on each chromosome were binned into 500 kb distance and the SNP with the lowest *P*-value was extracted from each bin. This step was performed to avoid fitting adjacent SNPs tagging the same QTL as fixed effects. Up to three most significant markers (FDR = 0.1) were selected separately for each fold in the five-fold cross-validation method. When no marker met this threshold, only the most significant marker was selected. The selected markers were then included in the GS+GWAS model as fixed effects while all the remaining markers from the full marker density were included as random effects. The GS+GWAS model was fitted in R (R Development Core Team 2016) using the ‘kinship.BLUP’ function in the rrBLUP package (Endelman 2011).

#### A model that accounts for genotype-by-environment interaction

The effect of modelling GEI on genomic prediction accuracy was assessed using a reaction norm model that incorporates the main effects and interactions of molecular markers and environments using covariance structures (EG-G×E) as described in Jarquín et al. (2014). The main effects of markers and environments were included in the model using techniques similar to the standard G-BLUP, while the interaction terms were modelled using a cell by cell product of two covariance structures, [Z_*g*_GZ′_*g*_]○[Z_*E*_Z′_*E*_], where Z_*g*_ is an incidence matrix for the vector of additive genetic effects, G is a marker-derived genomic relationship matrix, and Z_*E*_ represents the incidence matrix for the effects of environments (i.e., the matrix that connects the phenotypes with environments) (Jarquín et al. 2014). In this model, phenotypes (*y*_*ijk*_) are described as the sum of an overall mean (*μ*) plus a random deviation due to the environment (*E*_*i*_), which is a combination of site-years, plus marker covariates using marker-derived GRM (g_*j*_), plus an interaction term between genotypes and environments (*gE*_*ij*_) and a residual term (*ε*_*ij*_).

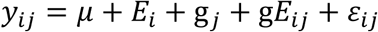

with *E*_*i*_ ∼ *N*(0, σ^2^_*E*_), g ∼ *N*(0, Gσ^2^_*g*_), gE ∼ *N* (0, [Z_*g*_GZ′_*g*_]○[Z_*E*_Z′_*E*_] σ^2^_*gE*_), ε_*ijk*_ ∼ *N*(0, σ^2^_ε_).

The accuracy of the model ‘EG-G×E’ that accounted for GEI was compared with a standard G-BLUP model plus a random environmental effect (*E*) yielding the ‘EG’ model without the interaction term which becomes

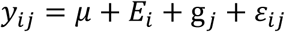

with *E*_*i*_ ∼ *N*(0, σ^2^_*E*_), g ∼ *N*(0, Gσ^2^_*g*_) and ε_*ijk*_ IID ∼ *N*(0, σ^2^_*ε*_).

Prediction accuracy of the reaction norm model was assessed using two cross-validation designs (CV1 and CV2) as implemented in Burgueño et al. (2012). Cross-validation one (CV1) involved predicting phenotypes of lines that have never been tested in any of the environments (newly developed lines). The second design (CV2) involved predicting phenotypes of lines that were evaluated in some environments but not in others (mimics incomplete field trials). Both CV1 and CV2 were implemented in a five-fold design. Prediction accuracy was computed as Pearson’s correlation between the predicted values and phenotypes of lines in the validation set within each environment and fold.

#### Multiple-trait prediction models

We used MT-BayesA (Jia and Jannink 2012) and MT-BayesA matrix (Jiang et al. 2015) models for joint prediction of multiple traits. Multiple-trait BayesA uses the genetic correlation between two or more traits to potentially improve the accuracy of prediction (Jia and Jannink 2012). Multiple-trait BayesA matrix is an antedependence-based model that uses correlations between traits as well as SNP effects simultaneously to improve prediction accuracy (Jiang et al. 2015). The standard GS models assume that marker effects are independently distributed; however, this is not always the case especially when adjacent SNPs are in high LD with the same QTL (Jiang et al. 2015). The antedependence model considers the potential nonstationary correlations between SNP effects near to a QTL (Yang and Tempelman 2012). MT-BayesA and MT-BayesA matrix models were fitted using C language programs developed by Jiang et al. (2015).

#### Genomic prediction scenarios

Three prediction scenarios were evaluated in this study: within-population, across-population, and across-environment. Within-population genomic predictions were made in each population using five-fold cross-validation design. In a five-fold cross-validation, each population was randomly divided into five groups of approximately equal sizes. In each fold, four groups were combined and used as the TP and predictions were made for the remaining group. This was repeated five times until predictions were made for each of the five groups. Across-population genomic predictions were made by using the LDP as TP to make predictions for LR-01 and LR-11 populations. LR-01 was used as TP to make prediction for LR-11 and vice versa. Across-environments predictions were made with the reaction norm model using CV1 and CV2 approaches described earlier. Accuracy of all models was determined as Pearson’s correlation (*r*) between the predicted values and observed phenotypes of lines in the validation set. The same cross-validation folds were used for single-trait, GS+GWAS, and multiple-trait models for a reliable comparison of their accuracy. Inferences for all Bayesian models were based on 30,000 iterations obtained after discarding 10,000 samples as a burn-in.

## Results

The distribution of all traits followed an approximately normal distribution in the LDP and LR-11 but it was slightly skewed towards higher values in LR-01 because some lines in LR-01 were susceptible to a residue of herbicide that had been applied the previous fall which led to delayed flowering and maturity (Online resource 1 to 3). There were strong positive correlations (*r* ≥ 0.72) among VEG, DTF, DTS, and DTM in the LDP and LR-11 populations (Online resource 1 and 2). Reproductive period was moderately correlated with DTM but showed weak correlations with the other traits in the LDP and LR-11. In LR-01, VEG was strongly correlated with DTF (*r* = 0.83) but weakly correlated with the other traits (Online resource 3). There was also a strong correlation among DTS, DTF, and REP in LR-01 (*r* ≥ 0.76) (Online resource 3). Broad-sense heritability was estimated using least-square means in each environment. A broad range of heritability estimates was obtained for all traits in each population (Table 1). The highest heritability was obtained for VEG in the LDP and LR-11, and REP in LR-01. Heritability of REP was the lowest in the LDP and LR-11. Trait heritability is one of the factors that affect GS accuracy, with prediction being more accurate for traits with higher heritability (Combs and Bernardo 2013; Heffner et al. 2011).

**Table 1.**
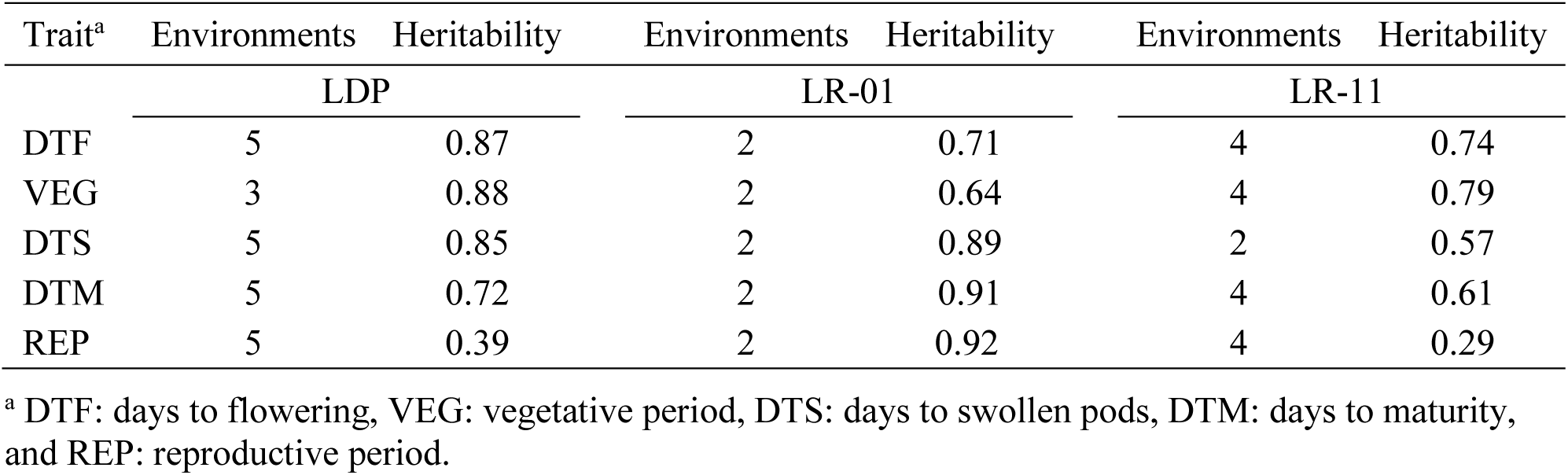
Broad-sense heritability estimates for five traits based on least square means from the respective number of environments in the LDP, LR-01, and LR-11 populations.

| Trait<sup>a</sup> | Environments | Heritability | Environments | Heritability | Environments | Heritability |
| --- | --- | --- | --- | --- | --- | --- |
|  | LDP |  | LR-01 |  | LR-11 |  |
| DTF | 5 | 0.87 | 2 | 0.71 | 4 | 0.74 |
| VEG | 3 | 0.88 | 2 | 0.64 | 4 | 0.79 |
| DTS | 5 | 0.85 | 2 | 0.89 | 2 | 0.57 |
| DTM | 5 | 0.72 | 2 | 0.91 | 4 | 0.61 |
| REP | 5 | 0.39 | 2 | 0.92 | 4 | 0.29 |

<sup>a</sup> DTF: days to flowering, VEG: vegetative period, DTS: days to swollen pods, DTM: days to maturity, and REP: reproductive period.

### Effect of population structure on prediction accuracy

The 324 lines in the LDP were clustered into three groups using marker-based K-means clustering. The first two principal components explained 33.5% of the genetic variance, and weakly differentiated the three groups indicating mild population structure (Fig. 1). The effect of population structure on model prediction accuracy was evaluated using an interaction model that partitioned marker effects into components that are constant across groups and interaction terms that describe group specific deviations (de los Campos et al. 2015).

**Fig. 1.**
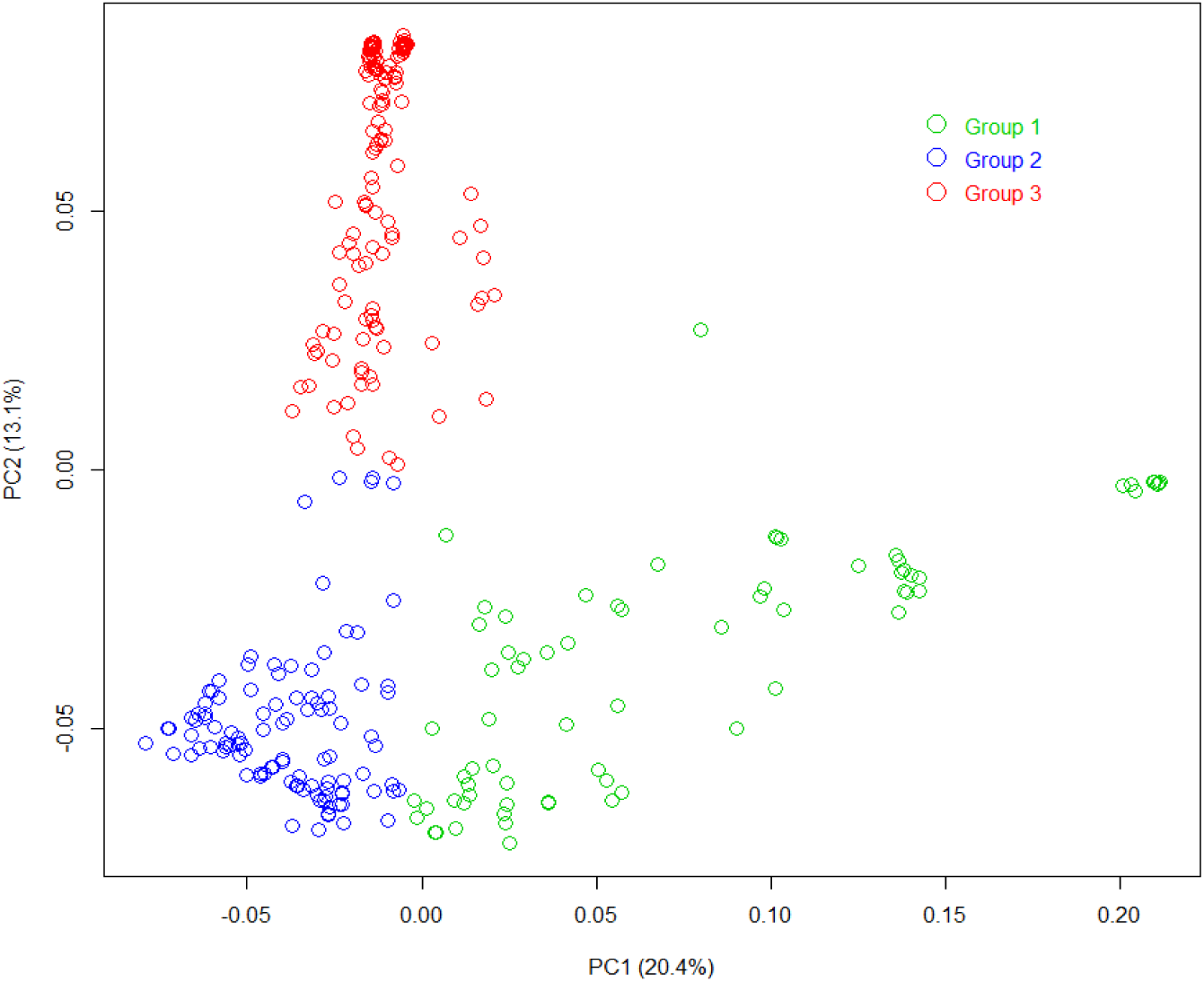
First two marker-derived principal components showing K-means clustering of the 324 lines in the LDP into three groups (Group one = 77 lines, group two = 106 lines, and group three = 141 lines).

Moderate to high prediction accuracies (ranging from 0.49 to 0.88) were obtained for all traits in the interaction and across-group models (Table 2). Accuracies varied among groups and traits, but it was consistently higher in group one followed by groups three and two, except for DTS where the accuracy in group two was slightly higher than in group three. Overall, the interaction model had similar accuracy to the across-group model that ignored population structure by assigning constant marker effects across groups (Table 2). This indicates that the effect of population structure on model prediction accuracy is negligible in this population.

**Table 2.** Average prediction accuracy (from 10 training-validation partitions) of the interaction and across-group models for five traits in the LDP.

| Trait<sup>a</sup> | Group<sup>b</sup> | Interaction model | Across-group model | % Change<sup>c</sup> |
| --- | --- | --- | --- | --- |
| DTF | 1 | 0.822 | 0.808 | 1.7 |
|  | 2 | 0.737 | 0.743 | -0.8 |
|  | 3 | 0.784 | 0.777 | 0.9 |
| VEG | 1 | 0.875 | 0.879 | -0.5 |
|  | 2 | 0.699 | 0.699 | 0.0 |
|  | 3 | 0.810 | 0.808 | 0.2 |
| DTM | 1 | 0.864 | 0.863 | 0.1 |
|  | 2 | 0.660 | 0.653 | 1.1 |
|  | 3 | 0.669 | 0.668 | 0.1 |
| DTS | 1 | 0.873 | 0.871 | 0.2 |
|  | 2 | 0.738 | 0.749 | -1.5 |
|  | 3 | 0.714 | 0.707 | 1.0 |
| REP | 1 | 0.734 | 0.709 | 3.5 |
|  | 2 | 0.487 | 0.494 | -1.4 |
|  | 3 | 0.758 | 0.757 | 0.1 |

<sup>a</sup> DTF: days to flowering, VEG: vegetative period, DTS: days to swollen pods, DTM: days to maturity, and REP: reproductive period.
<sup>b</sup> Groups identified from marker-based K-means clustering.
<sup>c</sup> Change in prediction accuracy of the interaction model relative to the prediction accuracy of the across-environment model.

### Within-population prediction accuracy

#### Single-trait prediction accuracy

Single-trait prediction accuracy varied across traits, regardless of the model, because they have different heritability estimates. Within-population prediction accuracies for all traits ranged from 0.65 to 0.85, 0.26 to 0.74, 0.41 to 0.79, and 0.28 to 0.59 in the LDP, LR-01, LR-11, and combined LR-01+LR-11 populations, respectively (Fig. 2). Prediction accuracy was lower for REP compared to the other traits in the LDP because heritability estimates were also lower (h^2^ = 0.39) and most of the phenotypic variance was caused by non-genetic factors. For each trait in the LDP, all models showed similar accuracy except for REP where the RKHS showed 3 to 5% higher accuracy than the other models. In LR-11, the accuracy of BayesA, BayesB, and GS+GWAS was higher for DTF and VEG (Fig. 2). GS+GWAS fitted three significant markers from fold specific single marker regression in the TP as fixed effects while all the remaining markers were fitted as random effects (Online resource 4). GS+GWAS showed higher accuracy compared to the standard RR-BLUP but its accuracy was either similar to or lower than that of BayesB (Fig. 2). Similarly, BayesB showed slightly higher accuracy for DTF and VEG in LR-01 and joint LR-01+LR-11 populations.

**Fig. 2.**
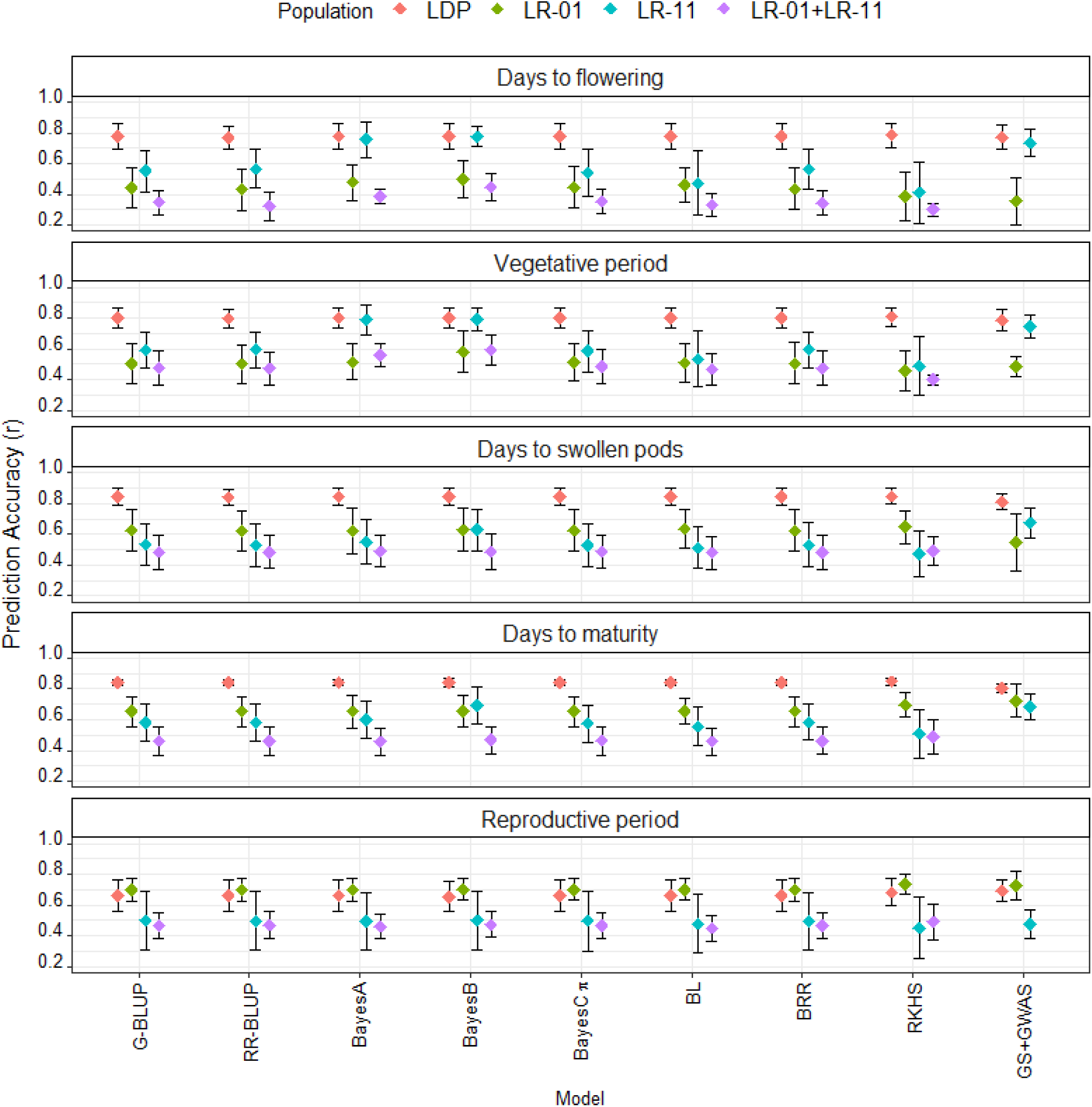
Average within-population prediction accuracy (based on five-fold cross-validation) of nine GS models for five traits measured in the lentil diversity panel (LDP), two RIL populations (LR-01 and LR-11) and a joint RIL population (LR-01 + LR-11). Vertical bars show standard deviation of the mean.

Prediction accuracy was high in the LDP compared to the biparental populations for all traits except for REP where the highest accuracy was obtained in LR-01, likely because its heritability estimate was also the highest in LR-01. Combining LR-01 and LR-11 populations resulted in either similar or lower accuracy compared to the accuracy within each population which suggests that there was no advantage of increasing TP size by combining the two biparental populations.

#### Multiple-trait prediction accuracy

We compared the accuracies of MT-BayesA and MT-BayesA matrix models with single-trait BayesA and the best single-trait model with the highest accuracy. In each population, two multiple-trait predictions were made using different trait combinations based on their phenotypic correlation (Table 3). In the LDP and LR-11, the first prediction was made for DTF, VEG, DTS, and DTM (*r* ≥ 0.72) while the second prediction was made for DTM and REP, correlations of *r* = 0.40 and 0.66 in the LDP and LR-11, respectively (Online resource 1 and 2). In LR-01, the first prediction was made for DTS, DTM, and REP (*r* ≥ 0.76) while the second prediction was made for DTF and VEG (*r* = 0.83) (Online resource 3). The accuracy of MT-GS models varied depending on the population and traits selected for joint prediction (Table 3). In the LDP, MT-BayesA and MT-BayesA matrix models showed 4 to 7% higher accuracy for REP compared to the accuracy of single-trait BayesA but accuracies remained the same for all other traits. MT-BayesA performed similar to the best single-trait model (RKHS) but MT-BayesA matrix showed 3% higher accuracy compared to the accuracy of RKHS for REP (Fig. 2 and Table 3). In LR-01, the accuracy of MT-GS models was either similar to or lower than the single-trait BayesA except for VEG when the joint prediction was made with DTF. The accuracy of MT-BayesA was 10% higher compared to the accuracy of single-trait BayesA for VEG but it was similar to the best single-trait model (BayesB) in LR-01. When joint prediction was made for DTM and REP in LR-11, MT-BatesA matrix resulted in 15 and 14% higher accuracy for REP compared to single-trait BayesA and the best single-trait model (BayesB), respectively (Table 3). Similarly, the accuracy of MT-GS models was higher than the accuracy of single-trait BayesA for DTM, but they had lower accuracy compared to BayesB in LR-11.

**Table 3.**
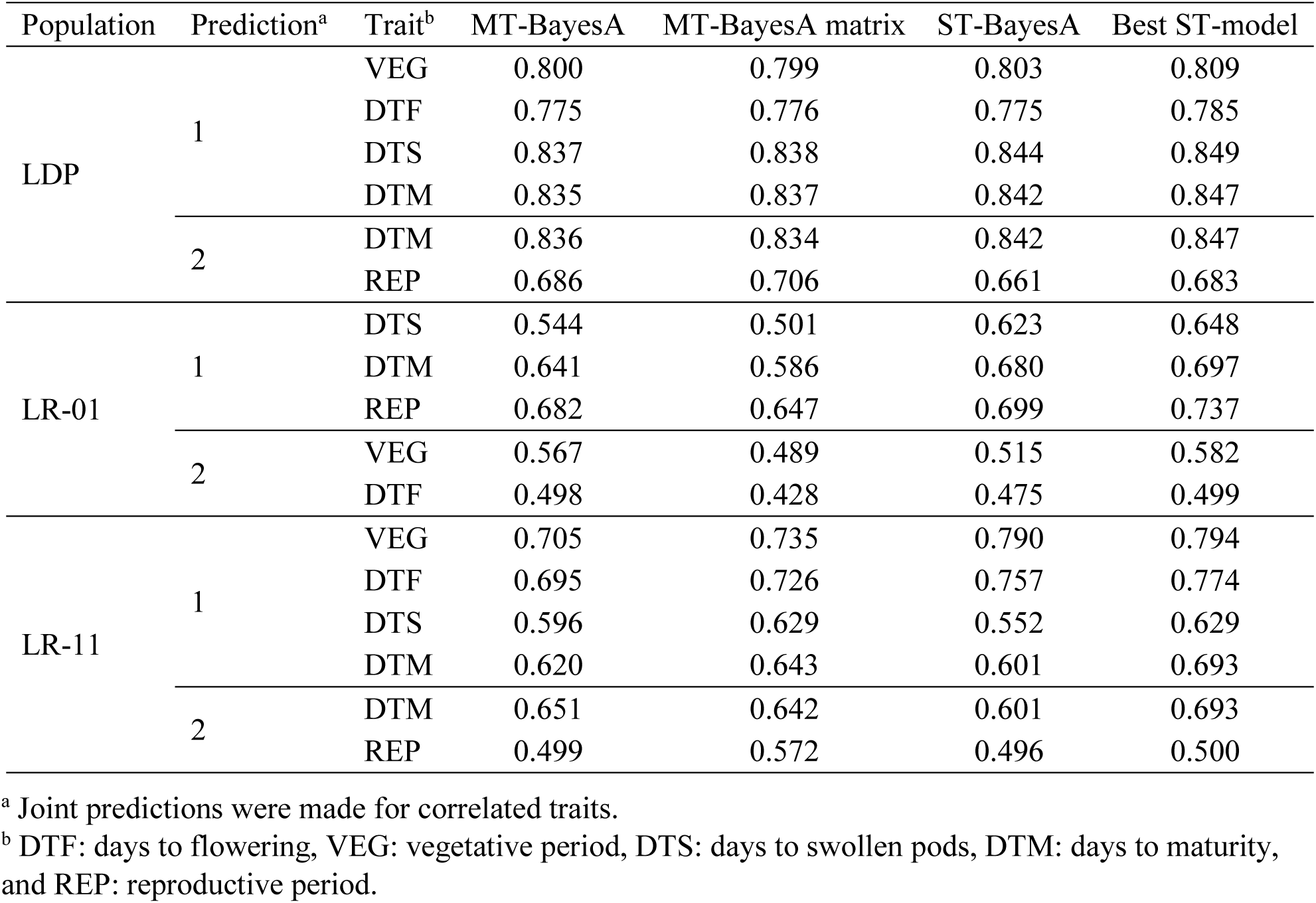
Average prediction accuracy (from five-fold cross-validation) of multiple-trait BayesA, multiple-trait BayesA matrix, single-trait BayesA, and the best single-trait prediction model in the LDP, LR-01, and LR-11 populations.

| Population | Prediction<sup>a</sup> | Trait<sup>b</sup> | MT-BayesA | MT-BayesA matrix | ST-BayesA | Best ST-model |
| --- | --- | --- | --- | --- | --- | --- |
| LDP | 1 | VEG | 0.800 | 0.799 | 0.803 | 0.809 |
|  |  | DTF | 0.775 | 0.776 | 0.775 | 0.785 |
|  |  | DTS | 0.837 | 0.838 | 0.844 | 0.849 |
|  |  | DTM | 0.835 | 0.837 | 0.842 | 0.847 |
|  | 2 | DTM | 0.836 | 0.834 | 0.842 | 0.847 |
|  |  | REP | 0.686 | 0.706 | 0.661 | 0.683 |
| LR-01 | 1 | DTS | 0.544 | 0.501 | 0.623 | 0.648 |
|  |  | DTM | 0.641 | 0.586 | 0.680 | 0.697 |
|  |  | REP | 0.682 | 0.647 | 0.699 | 0.737 |
|  | 2 | VEG | 0.567 | 0.489 | 0.515 | 0.582 |
|  |  | DTF | 0.498 | 0.428 | 0.475 | 0.499 |
| LR-11 | 1 | VEG | 0.705 | 0.735 | 0.790 | 0.794 |
|  |  | DTF | 0.695 | 0.726 | 0.757 | 0.774 |
|  |  | DTS | 0.596 | 0.629 | 0.552 | 0.629 |
|  |  | DTM | 0.620 | 0.643 | 0.601 | 0.693 |
|  | 2 | DTM | 0.651 | 0.642 | 0.601 | 0.693 |
|  |  | REP | 0.499 | 0.572 | 0.496 | 0.500 |

<sup>a</sup> Joint predictions were made for correlated traits.
<sup>b</sup> DTF: days to flowering, VEG: vegetative period, DTS: days to swollen pods, DTM: days to maturity, and REP: reproductive period.

#### Across-population prediction accuracy

Accuracies for across-population genomic prediction were very low for all traits relative to within-population prediction accuracy except for REP in LR-01 (Figs. 2 and 3). When the LDP was used as TP to make prediction in LR-01 and LR-11, accuracies for all traits ranged from to 0.57, and 0.09 to 0.41, respectively (Fig. 3a and b). In LR-11, accuracies were close to zero for DTF but relatively higher accuracies (ranging from 0.25 to 0.41) were obtained for the other traits (Fig. 3a). In LR-01, prediction accuracy was the highest for REP (ranged from 0.54 to 0.56) which also had the highest heritability (Fig. 3b). RKHS consistently resulted in higher across-population prediction accuracies for all traits when the LDP was used to make prediction in LR-01 and LR-11 (Figs. 3a and b). When one biparental population was used to make prediction for the other biparental population, accuracies were close to zero or negative for DTF and VEG in all models except in BayesA and BayesB (Fig. 3c and d). In contrast, slightly higher accuracies (ranging from 0.26 to 0.38) were obtained for DTS and DTM in LR-01 (Fig. 3c).

**Fig. 3.**
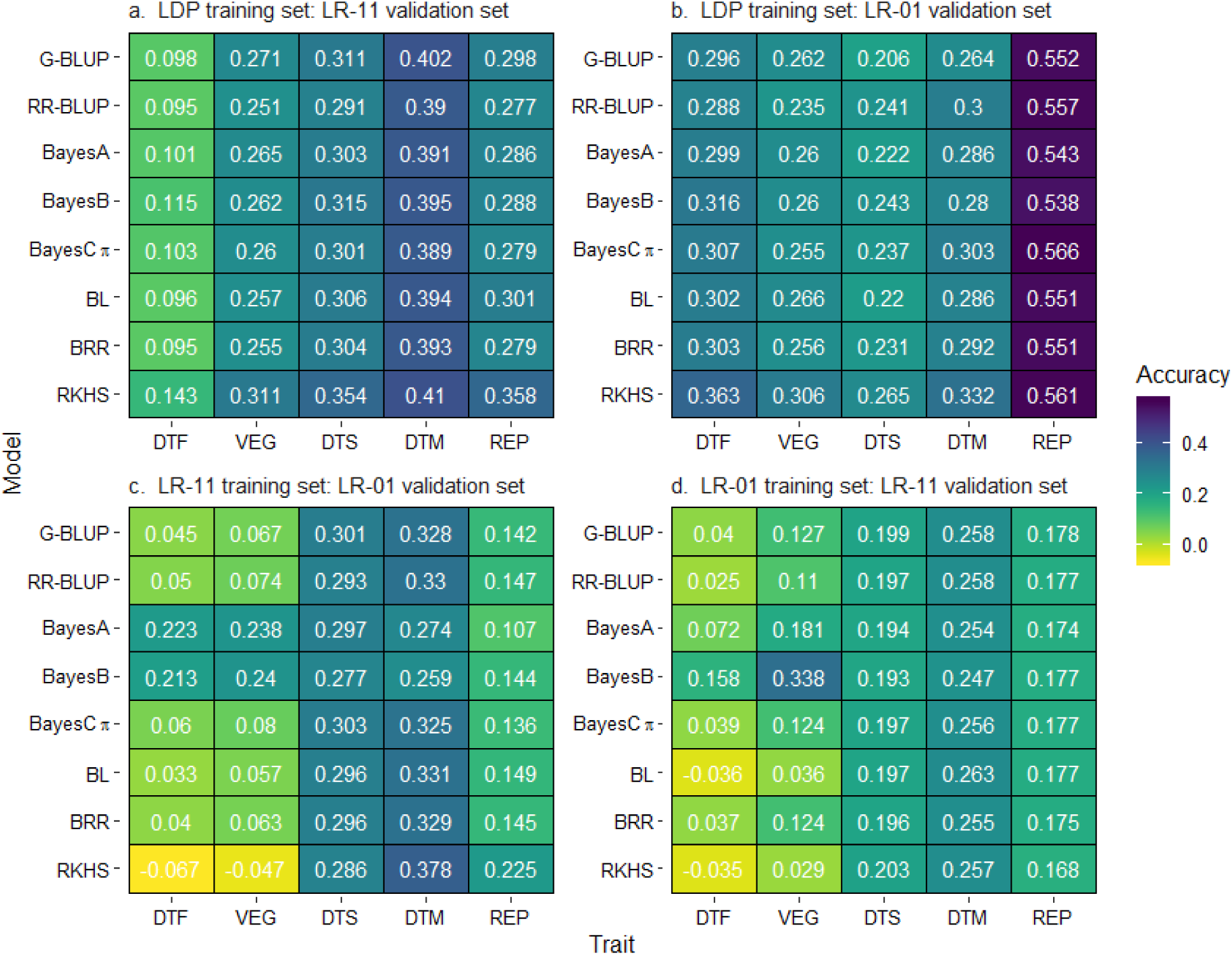
Across-population prediction accuracy of eight models for days to flowering (DTF), vegetative period (VEG), days to swollen pods (DTS), days to maturity (DTM), and reproductive period (REP).

#### Across-environment prediction accuracy

The effect of modelling GEI in GS was evaluated using two cross-validation designs that simulate prediction for newly developed lines (CV1) and incomplete field trials (CV2). Prediction accuracies varied depending on the cross-validation design and population. For all traits in the LDP, prediction accuracies ranged from 0.52 to 0.82 in CV1 and 0.63 to 0.95 in CV2 (Table 4). The lowest accuracies were obtained for REP in both cross-validation designs. Modelling GEI resulted in 7 to 30 and 4 to 35% higher accuracy for REP in CV1 and CV2, respectively (Table 4). Similarly, there was a 4 to 6% increase in prediction accuracy for DTM when modelling GEI in Sutherland 2016. But for all other traits, modelling the interaction term did not improve the accuracy of prediction compared to the model that included only the main effects of environments and markers. In LR-01, accuracies for all traits ranged from 0.41 to 0.66 and 0.63 to 0.88 in CV1 and CV2, respectively (Table 4). Both models showed similar accuracy for all traits in LR-01 indicating there was no benefit of modelling GEI when trait heritability is high. Similarly, across-environment prediction accuracies in LR-11 ranged from 0.19 to 0.60 and 0.22 to 0.89 for all traits in CV1 and CV2, respectively. The lowest accuracies were obtained for REP which also had low heritability (h^2^ = 0.29). Similar accuracies were obtained between the two models for all traits in LR-11 except for REP in CV2 where modelling GEI resulted in 18 to 66% higher accuracy. This shows that modelling the GEI term improved the accuracy of prediction for a low heritability trait but modelling only the main effects was enough for traits of high heritability. Overall, CV2 consistently resulted in higher accuracy compared to CV1 for all traits and populations because CV2 allows borrowing of information for the same line across environments (Table 4).

**Table 4.** Average across-environment prediction accuracy (from five-fold design) based on Pearson’s correlation between predicted and actual phenotypes in each environment. Predictions were made using two models and cross-validation designs (CV1 and CV2).

| Population | Environment | DTF<sup>a</sup> | DTM | DTS | REP | DTF | DTM | DTS | REP |
| --- | --- | --- | --- | --- | --- | --- | --- | --- | --- |
|  |  | Model EG |  |  |  | Model EG-G×E |  |  |  |
|  |  | CV1<sup>b</sup> |  |  |  |  |  |  |  |
| LDP | Rosthern 2016 | 0.749 | 0.766 | 0.797 | 0.488 | 0.734 | 0.761 | 0.802 | 0.524 |
|  | Rosthern 2017 | 0.802 | 0.738 | 0.794 | 0.558 | 0.803 | 0.752 | 0.798 | 0.597 |
|  | Sutherland 2016 | 0.798 | 0.746 | 0.807 | 0.529 | 0.796 | 0.777 | 0.811 | 0.609 |
|  | Sutherland 2017 | 0.812 | 0.769 | 0.822 | 0.490 | 0.819 | 0.779 | 0.822 | 0.638 |
|  | Sutherland 2018 | 0.787 | 0.729 | 0.806 | 0.585 | 0.785 | 0.746 | 0.821 | 0.699 |
|  | CV2 |  |  |  |  |  |  |  |  |
|  | Rosthern 2016 | 0.896 | 0.856 | 0.875 | 0.595 | 0.901 | 0.861 | 0.877 | 0.632 |
|  | Rosthern 2017 | 0.943 | 0.826 | 0.890 | 0.625 | 0.947 | 0.840 | 0.915 | 0.649 |
|  | Sutherland 2016 | 0.950 | 0.811 | 0.922 | 0.579 | 0.953 | 0.857 | 0.921 | 0.698 |
|  | Sutherland 2017 | 0.958 | 0.864 | 0.933 | 0.558 | 0.950 | 0.878 | 0.936 | 0.720 |
|  | Sutherland 2018 | 0.923 | 0.807 | 0.903 | 0.526 | 0.931 | 0.812 | 0.925 | 0.708 |
| LR-01 | CV1 |  |  |  |  |  |  |  |  |
|  | Rosthern 2018 | 0.484 | 0.573 | 0.543 | 0.608 | 0.485 | 0.576 | 0.548 | 0.611 |
|  | Sutherland 2018 | 0.408 | 0.613 | 0.552 | 0.654 | 0.406 | 0.615 | 0.559 | 0.656 |
|  | CV2 |  |  |  |  |  |  |  |  |
|  | Rosthern 2018 | 0.637 | 0.831 | 0.836 | 0.849 | 0.638 | 0.850 | 0.837 | 0.854 |
|  | Sutherland 2018 | 0.631 | 0.870 | 0.765 | 0.882 | 0.630 | 0.862 | 0.767 | 0.867 |
| LR-11 | CV1 |  |  |  |  |  |  |  |  |
|  | Rosthern 2017 | 0.587 | 0.499 | – | 0.295 | 0.585 | 0.499 | – | 0.299 |
|  | Rosthern 2018 | 0.457 | 0.331 | 0.428 | 0.296 | 0.456 | 0.331 | 0.430 | 0.299 |
|  | Sutherland 2017 | 0.595 | 0.575 | – | 0.194 | 0.595 | 0.574 | – | 0.192 |
|  | Sutherland 2018 | 0.512 | 0.507 | 0.440 | 0.393 | 0.514 | 0.509 | 0.442 | 0.395 |
|  | CV2 |  |  |  |  |  |  |  |  |
|  | Rosthern 2017 | 0.891 | 0.738 | – | 0.370 | 0.890 | 0.736 | – | 0.521 |
|  | Rosthern 2018 | 0.736 | 0.545 | 0.597 | 0.383 | 0.736 | 0.546 | 0.597 | 0.453 |
|  | Sutherland 2017 | 0.889 | 0.794 | – | 0.220 | 0.889 | 0.792 | – | 0.364 |
|  | Sutherland 2018 | 0.892 | 0.770 | 0.640 | 0.326 | 0.892 | 0.769 | 0.639 | 0.442 |

<sup>a</sup> DTF: days to flowering, VEG: vegetative period, DTS: days to swollen pods, DTM: days to maturity, and REP: reproductive period.
<sup>b</sup> CV1: prediction for lines that were not evaluated in any of the environments, CV2: prediction for lines that were evaluated in some of the environments.

## Discussion

This study evaluated the accuracy of single-trait models, multiple-trait models, a model that included significant markers from GWAS as fixed effects, and models that accounted for population structure and GEI using a lentil diversity panel, two RIL populations, and a joint RIL population. Moreover, within-population, across-population, and across-environment prediction were tested to simulate scenarios that breeders may face when implementing GS. Highly variable prediction accuracies were obtained depending on the trait, population, prediction scenario, and statistical model used.

### Effect of population structure on genomic prediction accuracy

We evaluated the effect of population structure on genomic prediction accuracy in the LDP. Population structure is known to cause spurious associations in GWAS (Pritchard and Rosenberg 1999). Several studies have also shown that genomic predictions can be biased in the presence of a strong population structure (Asoro et al. 2011; Isidro et al. 2015; Saatchi et al. 2011). We compared the accuracies of an interaction model that accounted for population structure with an across-group model that ignored population structure by considering constant marker effects across sub-populations (de los Campos et al. 2015). The interaction model resulted in similar accuracy to the across-group model indicating no effect of population structure on genomic prediction accuracy (Table 2). The effect of population structure on genomic prediction accuracy varies depending on prediction strategies, genetic architectures of traits and populations (Guo et al. 2014). de los Campos et al. (2015) compared the interaction model with an across-group and within-group models using a wheat data set with strong population structure and a pig data set with less marked differentiations between sub-populations. The interaction model performed slightly better or similar to the best performing model (either across-group or within-group analysis) in both data sets. In contrast, Crossa et al. (2016) indicated that prediction accuracy decreased by 15-20% compared to the accuracy obtained without correcting for population structure in wheat landraces. Similarly, Guo et al. (2014) showed that accounting for population structure reduced the accuracy of genomic predictions when there is population structure in both the training and validation sets. This indicates that the benefit of accounting for population structure in GS may vary in different populations and it needs to be assessed on a case by case basis.

### Within-population genomic prediction

Nine single-trait and two MT-GS models were used to make within-population genomic predictions based on a five-fold cross-validation design. The main difference among the evaluated single-trait models is in their assumptions about the variances of marker effects. RR-BLUP includes all markers as random factors and uses a penalty term that shrinks their effects uniformly towards zero because all loci are assumed to explain equal amount of the genetic variance (Meuwissen et al. 2001; Whittaker et al. 2000). G-BLUP has similar assumption to RR-BLUP but the genomic relationship matrix, which estimates the realized proportion of the genome that is shared by two individuals, is used to estimate marker effects (Habier et al. 2013; VanRaden 2008). Fitting all markers as random effects and shrinking their effects uniformly has disadvantages because this method fails to distinguish between markers tagging known major genes and random genome-wide markers (Bernardo 2014). GS+GWAS combines RR-BLUP with significant markers from GWAS in the TP fit as fixed effects (Spindel et al. 2016). In Bayesian models, the variance explained by each locus is allowed to vary and depends on prior assumptions about the distributions of marker effects, which determines the extent and type of shrinkage (de los Campos et al. 2013; Meuwissen et al. 2001).

Within-population prediction accuracies ranged from 0.65 to 0.85, 0.26 to 0.74, 0.41 to 0.79, and 0.28 to 0.59 for all traits in the LDP, LR-01, LR-11, and combined LR-01+LR-11 populations, respectively (Fig. 2). Accuracies were generally higher in the LDP compared to the biparental populations because the number of lines in the LDP is nearly three times that of the biparental populations. Training population size is an important factor that affects the accuracy of genomic predictions. Previous studies showed that increasing the TP size increases the accuracy of prediction because it provides more data to estimate marker effects (Asoro et al. 2011; Saatchi et al. 2010; VanRaden et al. 2009). All models showed similar accuracy in the LDP except for RKHS that showed 3 to 5% higher accuracy than the other models for REP. BayesA, BayesB, and GS+GWAS resulted in higher accuracy for DTF and VEG in LR-11 (Fig. 2). In theory, RR-BLUP is expected to have higher accuracy for traits controlled by many QTL with small effects, while Bayesian models have higher accuracy when few QTL with large effects control most of the phenotypic variance (Lorenz et al. 2011). The higher accuracy of BayesA and BayesB in LR-11 could be due to the presence of large effect QTL controlling DTF and VEG.

Similar accuracies were obtained between MT-GS and single-trait models when joint predictions were made for correlated high heritability traits (Tables 1 and 3). However, the accuracy of MT-BayesA matrix increased by 3 and 14% for REP when it was predicted with DTM in the LDP and LR-11, respectively (Table 3). Jiang et al. (2015) also showed that the prediction accuracy of the simplified version of MT-BayesA matrix model was 2.8 to 8.6% higher compared to the accuracy of the best single-trait prediction model. Previous studies based on simulated data showed that multiple-trait models have higher prediction accuracy for a low heritability trait (h^2^ = 0.1) jointly predicted with a correlated high heritability trait (h^2^ ≥ 0.5) (Hayashi and Iwata 2013; Jia and Jannink 2012; Jiang et al. 2015). However, no improvement in accuracy was observed compared to single-trait models for traits of high heritability and in the absence of genetic correlation between traits (Hayashi and Iwata 2013; Jia and Jannink 2012; Jiang et al. 2015).

RKHS regression uses a kernel function to convert the marker data into a square matrix capturing complex interactions potentially arising in whole-genome models (Gianola and van Kaam 2008). Unlike the other GS models that predict GEBVs (additive genetic effects), RKHS regression captures both the additive and non-additive genetic effects and predicts the total genetic value of individuals (Heslot et al. 2012). The accuracy of RKHS was slightly higher for REP in the LDP but it resulted in similar or lower accuracy than the other models for all traits (Fig. 2). This suggests that the contribution of non-additive genetic effects to the total genetic variance is negligible. Previous studies in wheat and maize also reported no benefit of models accounting for non-additive effects over simple additive models (Lorenzana and Bernardo 2009; Sallam et al. 2015; Zhao et al. 2013). In contrast, improved prediction accuracies were reported using RKHS and models that incorporate epistasis (Crossa et al. 2010; Pérez-Rodríguez et al. 2012). This suggests that the benefit of accounting for non-additive effects in GS may vary depending on the trait and population type.

GS+GWAS fitted up to three markers that are significantly associated with traits as fixed effects (Online resource 4). There was no benefit of fitting significant markers as fixed effects in the LDP because all markers explained less than 5% of the phenotypic variance (Online resource 4). GS+GWAS works best for traits with one or more medium to large effect QTL segregating in the population but it has no advantage if a trait has no significant GWAS peaks (Spindel et al. 2016). Bernardo (2014) suggested that major QTL should be fitted as having fixed effects in GS, especially if a few major QTL are present and if each QTL explains more than 10% of the genetic variance. Including up to three significant markers as a fixed effect in GS+GWAS resulted in 18 to 30% higher accuracy compared to the standard RR-BLUP for DTF, VEG, DTS and DTM in LR-11 (Fig. 2). Similarly, 4 and 11% higher accuracy was obtained for REP and DTM, respectively in LR-01. However, no improvement in accuracy was observed for the other traits in LR-01 and LR-11. Spindel et al. (2016) showed that GS+GWAS performed better than the standard RR-BLUP in all cases and up to 30% higher accuracy was obtained for flowering time of rice which had a large GWAS peak. Although GS+GWAS gave higher accuracies compared to the standard RR-BLUP in LR-01 and LR-11, its accuracy was either similar or lower to BayesB indicating that variable selection models such as BayesB can be equally effective to account for large effect QTL in GS.

Combining LR-01 and LR-11 to increase the TP size did not improve the accuracy of prediction for all traits (Fig. 2). Previous studies in wheat and barley also indicated that prediction accuracies did not improve when unrelated populations from different breeding programs were merged to increase TP size (Charmet et al. 2014; Lorenz et al. 2012). Unrelated populations may have different LD phase and combining them into a single TP reduces the overall LD (Goddard 2012). Moreover, combining multiple populations may create strong population structure and allele frequency differences between sub-populations which reduces GS accuracy (Riedelsheimer et al. 2013). The standard GS models assume that marker effects are constant across sub-populations and fail to account for differences in allele frequency due to population structure. Therefore, models that account for population structure need to be considered when different populations are combined to increase TP size.

### Across-population genomic prediction

Variable across-population prediction accuracies were obtained depending on the trait and population used as TP. Overall, across-population prediction accuracies were very low relative to within-population prediction accuracy for all traits except for REP in LR-01 (Figs. 2 and 3). The prediction accuracy was the highest (ranged from 0.54 to 0.56) for REP which indicates that moderate across-population prediction can be made for highly heritable traits (Fig. 3b). Prediction in LR-01 and LR-11 ranged from 0.21 to 0.57 and 0.09 to 0.41, respectively when the LDP was used as TP (Fig. 3a and b). When one biparental population was used to predict the other biparental population, accuracies were close to zero or negative for DTF and VEG in all models except in BayesA and BayesB (Fig. 3c and d). Previous studies in wheat and maize also reported mean accuracies of zero or negative values when unrelated TP is used to make prediction for an independent population (Charmet et al. 2014; Crossa et al. 2014; Riedelsheimer et al. 2013; Windhausen et al. 2012). GS models utilize both the genetic relationships among individuals and LD between markers and QTL to improve prediction accuracy (Habier et al. 2007). Across-population GS prediction not only requires strong LD but similar linkage phases in each population (Goddard and Hayes 2007). When the TP is unrelated to the breeding population, marker effects estimated in one population cannot be transferred to the other population due to differences in allele frequency and LD phase resulting in low prediction accuracy (Bassi et al. 2016; Windhausen et al. 2012). In plant breeding, early generation nursery is comprised of different families from many crosses and the most attractive application of GS is to estimate GEBVs of lines in these families based on marker effects estimated from an independent population. However, the results of this study, as well as previous studies, showed that GS has very low accuracy for such an application. Based on a simulation study, Meuwissen (2009) suggested that a substantially higher marker density and TP size is required for accurate prediction of GEBVs in unrelated individuals. Therefore, across-population prediction should not be considered unless the two populations are closely related, or the size of TP is very large.

### Across-environment genomic prediction

Across-environment predictions were made using reaction norm models that incorporated the main and interaction terms of environments and markers (Jarquín et al. 2014). Genotype-by-environment interaction is an important factor that affects both phenotypic and genomic selection accuracy. Across-environment prediction accuracies in CV1 ranged from 0.49 to 0.82, 0.41 to 0.66, and 0.19 to 0.60 in the LDP, LR-01, and LR-11, respectively (Table 4). Prediction accuracies in CV2 were consistently higher compared to CV1 and ranged from 0.53 to 0.96, 0.63 to 0.88, and 0.22 to 0.89 in the LDP, LR-01, and LR-11, respectively. Similar results were obtained in previous studies that used similar cross-validation designs (Burgueño et al. 2012; Crossa et al. 2015; Jarquín et al. 2014; Jarquín et al. 2017; Lopez-Cruz et al. 2015; Pérez-Rodríguez et al. 2015). Modelling the interaction term improved the accuracy of prediction for REP by 4 to 35% and 18 to 66% in the LDP and LR-11, respectively (Table 4). But for all other traits modelling GEI resulted in similar accuracies with the model that included only the main effects of markers and environments. Previous studies showed that modelling GEI improves the accuracy of genomic predictions (Crossa et al. 2015; Cuevas et al. 2017; Jarquín et al. 2014; Jarquín et al. 2017; Lopez-Cruz et al. 2015). Jarquín et al. (2014) used the reaction norm model to predict grain yield of wheat and reported a 35% increase in accuracy when adding interaction terms between markers and environments. Recently, Jarquín et al. (2017) showed that modelling GEI resulted in 16 to 82% higher accuracy than a baseline model which did not include the GEI term. Other studies in wheat and cotton that used the reaction norm model with pedigrees instead of molecular markers also obtained the highest prediction accuracy when GEI term was included in the model (Pérez-Rodríguez et al. 2015; Sukumaran et al. 2017).

## Conclusion

This research provides the first empirical evidence of GS in lentil. Comparison of different GS models and approaches showed that most single-trait GS models have similar accuracy in the absence of large effect QTL underlying traits but BayesB is the best model when QTL with relatively large effects are present. GS models that accounted for GEI and multiple-trait models improved the accuracy of prediction for a low heritability trait, but they have no advantage when trait heritability is high. Within-population and across-environment genomic predictions resulted in moderate to high accuracies, but across-population prediction accuracies were very low. This suggests that GS can be implemented in lentil to make within-population prediction using the parental generation of a cross as TP to predict GEBVs of progenies or across-environment predictions in incomplete field trials, but across-population prediction should not be considered when the population size is small.

## Supporting information

Online resource 1

Online resource 2

Online resource 3

Online resource 4

## Acknowledgements

This research was conducted as part of the “Application of Genomics to Innovation in the Lentil Economy (AGILE)”, project funded by Genome Canada and managed by Genome Prairie. We are grateful for the matching financial support from the Saskatchewan Pulse Growers, Western Grains Research Foundation, the Government of Saskatchewan, and the University of Saskatchewan. We acknowledge the technical assistance of the bioinformatics, field and molecular lab staff of the Pulse Crop Breeding and Genetics group at the University of Saskatchewan. We thank Ms. Crystal Chan, AGILE project coordinator, for her assistance during this research.

## Author contribution statement

TAH generated phenotypic data, performed all statistical analyses, and wrote the manuscript. TH, DW, SN generated phenotypic data, and edited the manuscript. LR generated the SNP data set. AV designed the manuscript and provided germplasm materials. KEB designed the experiment, supervised the project, and edited the manuscript. All authors read and approved the final manuscript.

