## Supplementary material for "Genomic selection for lentil breeding: empirical evidence": Online resource 1

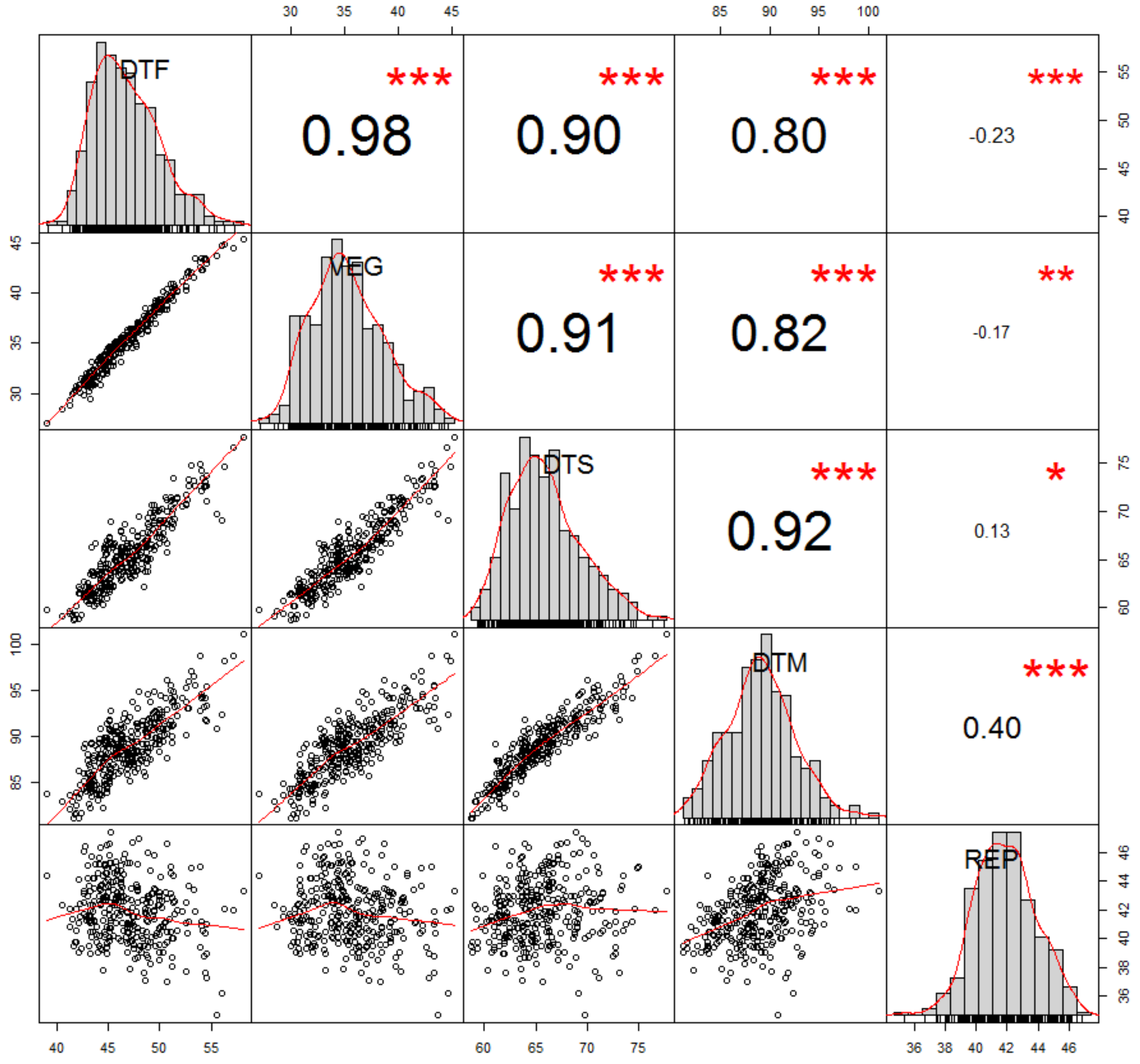


**Online Resource 1** Frequency distribution and pairwise correlations for days to flowering (DTF), vegetative period (VEG), days to swollen pods (DTS), days to maturity (DTM), and reproductive period (REP) in the LDP.
